# Decoding Single-Cell Omics of Perturbation Responses Using DeSCOPE

**DOI:** 10.64898/2026.04.13.718147

**Authors:** Pengpeng Wu, Hailin Wei, Yazi Li, Xinwei Zheng, Caibin Zhou, Xihao Hu, Chenfei Wang

## Abstract

Deciphering cellular responses to genetic perturbations is fundamental to modeling gene regulatory networks and understanding mechanisms that change cellular phenotypes. However, current computational approaches often fail to outperform simple baseline models, highlighting a critical bottleneck in their generalizability and robustness. Here, we present DeSCOPE, a lightweight conditional variational autoencoder framework for predicting genetic perturbation responses spanning transcriptomic, epigenomic, and broader multi-modal landscapes. We systematically benchmarked DeSCOPE across diverse datasets under two challenging out-of-distribution settings: unseen genes and unseen cell types. DeSCOPE uniquely surpasses simple baselines in the unseen gene scenario, and achieves substantially improved performance for unseen cell types while requiring fine-tuning with far fewer perturbed genes. Finally, DeSCOPE demonstrates superior performance in predicting combinatorial multi-gene perturbations. Overall, DeSCOPE serves as a versatile multi-modal virtual cell model that can effectively guide the design of therapeutic targets that change cellular phenotypes. DeSCOPE is available at https://github.com/wanglabtongji/DeSCOPE.

## Introduction

Elucidating how cells respond to genetic perturbations is essential for understanding the mechanisms of cellular phenotype regulation^1^. The rapid development of single-cell resolution technologies has facilitated systematic, multi-dimensional characterization of these responses at transcriptomic^2, 3, 4^, epigenomic^5, 6, 7^, proteomic^8, 9^, and morphological^10^ levels. These advances have laid a solid foundation for developing virtual cell models, a longstanding goal in computational biology^11, 12, 13^. Virtual cell models trained on single-cell perturbation data enable in silico prediction of cellular responses to genetic perturbations^14^. Building upon this foundation, this capability holds transformative potential in biomedical research and applications. Through systematic in silico perturbation of cellular systems, these models can identify causal drivers of phenotypic changes, thereby accelerating target discovery and early-stage therapeutic development^15, 16, 17, 18, 19^. Moreover, they provide a powerful framework for dissecting gene regulatory networks underlying cellular phenotype changes and for rationally manipulating cellular trajectories^20, 21^. In the long term, virtual cell models hold the promise of realizing individualized digital twins for dynamic health monitoring and precision therapeutic interventions^22, 23^.

Although high-throughput CRISPR screening has matured considerably, scaling these assays remains prohibitively costly and time-consuming^24^, spurring the development of computational methods to circumvent experimental bottlenecks. Recent advances include GEARS^25^, which encodes Gene Ontology (GO) priors into gene interaction graphs to decipher perturbation-induced transcriptomic responses via graph neural networks. In parallel, STATE^26^ adopts a population-level modeling paradigm, treating individual cells as tokens to capture relationships among cell populations through attention mechanisms. Furthermore, emerging foundation models such as scGPT^27^, scFoundation^28^, and EpiAgent^29^ distill universal biological knowledge from vast wild-type atlases during pretraining, enabling precise perturbation prediction with efficient fine-tuning. Nevertheless, the effectiveness of these models has been increasingly scrutinized by recent research^30, 31, 32^. Evidence reveals that these sophisticated approaches frequently yield no improvement over rudimentary linear models, occasionally falling short of even a simple mean baseline. Such observations underscore the substantial hurdles that remain in advancing these computational methodologies. Moreover, most existing methods are predominantly tailored to transcriptomic data, often exhibiting limited generalizability or failing outright when extended to other modalities such as proteomics or epigenomics. Consequently, constructing a universal single-cell perturbation model capable of robust cross-modal application has become a critical imperative.

To overcome the limitations of existing approaches, we propose DeSCOPE, a lightweight and efficient virtual cell model for accurate prediction of single-cell responses to genetic perturbations. DeSCOPE leverages gene embeddings derived from the protein language model ESM2^33^ and explicitly decouples the latent distributions of control and perturbed cells, enabling robust out-of-distribution generalization across two demanding scenarios^34^: unseen genes and unseen cell types. Across multiple benchmark datasets, DeSCOPE consistently demonstrates superior performance and outperforms other baseline models. In addition, we benchmarked DeSCOPE on challenging generalization settings, including double-gene perturbation prediction and epigenome prediction. Our results establish DeSCOPE as a robust and efficient general-purpose framework for predicting gene perturbation effects across complex biological contexts.

## Results

### Overview of DeSCOPE

DeSCOPE is a lightweight and scalable virtual cell model built upon a conditional variational autoencoder (cVAE) architecture (Fig. 1). Given unperturbed cellular states represented by a feature matrix from gene expression (or other modalities) and target perturbation genes as inputs, the model predicts post-perturbation cellular states. DeSCOPE leverages the protein language model ESM2 to derive gene-specific embeddings, enabling structured representation of heterogeneous perturbation responses. These embeddings serve as conditioning variables that guide the encoder to learn prior and posterior distributions within perturbation-aware latent subspaces. During decoding, the embeddings are combined with sampled latent variables to reconstruct cellular states characterized by distinct perturbation signatures (Extended Data Fig. 1).

**Fig. 1.**
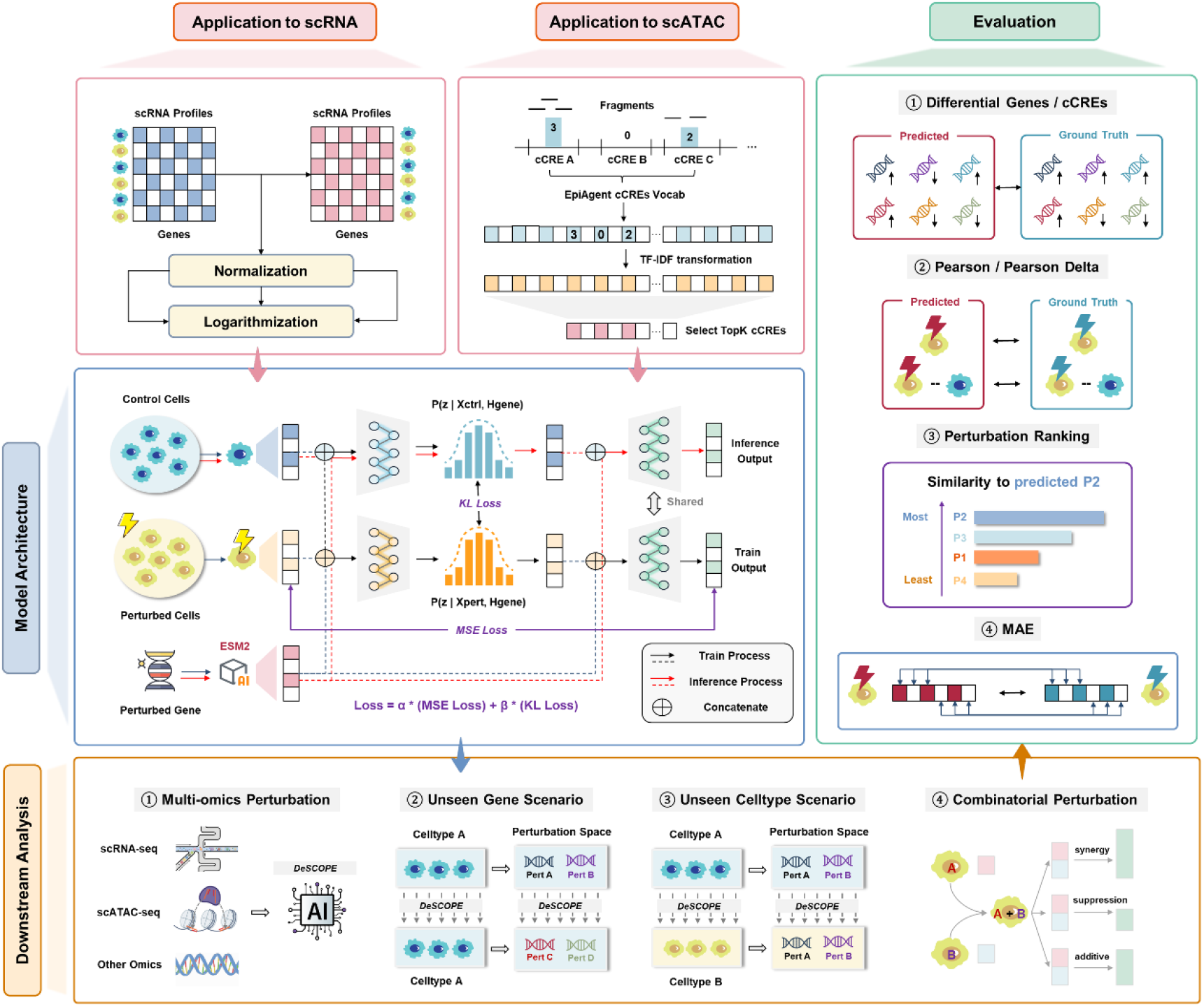
Overview of DeSCOPE. DeSCOPE takes scRNA-seq or scATAC-seq data as input. For scRNA-seq, gene expression matrices are normalized and log-transformed prior to modeling. For scATAC-seq, peaks are mapped to cis-regulatory elements (cCREs) using the EpiAgent vocabulary, followed by term frequency-inverse document frequency (TF-IDF) transformation and selection of the top k regulatory elements. The framework is implemented as a cVAE, in which perturbation gene embeddings derived from ESM2 are incorporated as conditioning variables to align latent distributions of control and perturbed cells via a KL divergence objective, while the decoder reconstructs perturbation-specific cellular states using a mean squared error (MSE) loss. Model outputs are evaluated using complementary metrics, including differential gene or cCRE prediction, Pearson correlation, perturbation ranking consistency, and mean absolute error (MAE).

To promote biological coherence, DeSCOPE incorporates regularization constraints that encourage alignment between the prior distribution inferred from control cells and the posterior distribution derived from perturbed cells. This design reflects the empirical observation that perturbed cellular states typically remain proximal to their matched controls, exhibiting localized manifold shifts rather than global restructuring. The resulting regularization preserves topological continuity in latent space and enhances generalization under sparse training conditions and unseen perturbation combinations.

A defining strength of DeSCOPE is its scalability across molecular modalities. By parameterizing both the encoder and decoder as lightweight multilayer perceptrons (MLPs), the framework substantially reduces computational complexity while maintaining the capacity to model extremely high-dimensional inputs. DeSCOPE captures long-range chromatin accessibility patterns in scATAC-seq perturbation datasets and supports modeling of gene expression programs spanning large subsets of genes to transcriptome-wide scales in scRNA-seq perturbation experiments. This design mitigates architectural constraints associated with input dimensionality and sequence length in conventional approaches. Furthermore, the framework is intrinsically extensible to emerging perturbation modalities, including spatial transcriptomics and integrative multiomics profiling, providing a generalizable computational basis for constructing comprehensive virtual cell perturbation atlases.

### Generalization to unseen genes in transcriptomic perturbation prediction

To comprehensively benchmark DeSCOPE, we collected five single-gene perturbation scRNA-seq datasets from different cell lines, including K562 (K562_ESSENTIAL), RPE1, H1, HepG2, and Jurkat. We evaluated DeSCOPE’s performance in predicting transcriptional responses to unseen gene perturbations and compared it with four deep-learning-based methods, including STATE, CPA^35^, scGPT, GEARS, and two simple baseline methods, including Linear and PerturbMean. Because scGPT requires perturbed genes to be a subset of detected genes and GEARS requires perturbed genes to appear in the prior Gene Ontology (GO) graphs, we retained only the overlapping perturbations for downstream benchmarking. Model performance was evaluated using several complementaryt metrics, including the overlap ratio of differentially expressed (DE) genes (Overlap@50 and Overlap@100), Pearson correlation between predicted and true expression changes across all genes (Pearson Δ) or across the top 20 DE genes (Pearson Δ20), mean absolute error of all genes (MAE), and a perturbation discrimination score (PDS) that quantifies whether predicted perturbation effects remain distinguishable.

Across the five datasets, DeSCOPE consistently achieved superior performance across these evaluation metrics (Fig. 2a and Extended Data Fig. 2a). In particular, DeSCOPE obtained the highest Overlap@50, Overlap@100, and PDS values across all benchmarked datasets, highlighting its strong ability to accurately capture perturbation-induced differential expression patterns. Meanwhile, DeSCOPE also achieved competitive MAE, Pearson Δ, and Pearson Δ20 scores compared with other methods, indicating improved accuracy in predicting gene expression responses to unseen perturbations. In addition, a simple runtime comparison with several representative methods shows that DeSCOPE requires less training time and GPU memory while maintaining strong predictive performance (Extended Data Table 1, Methods).

**Fig. 2.**
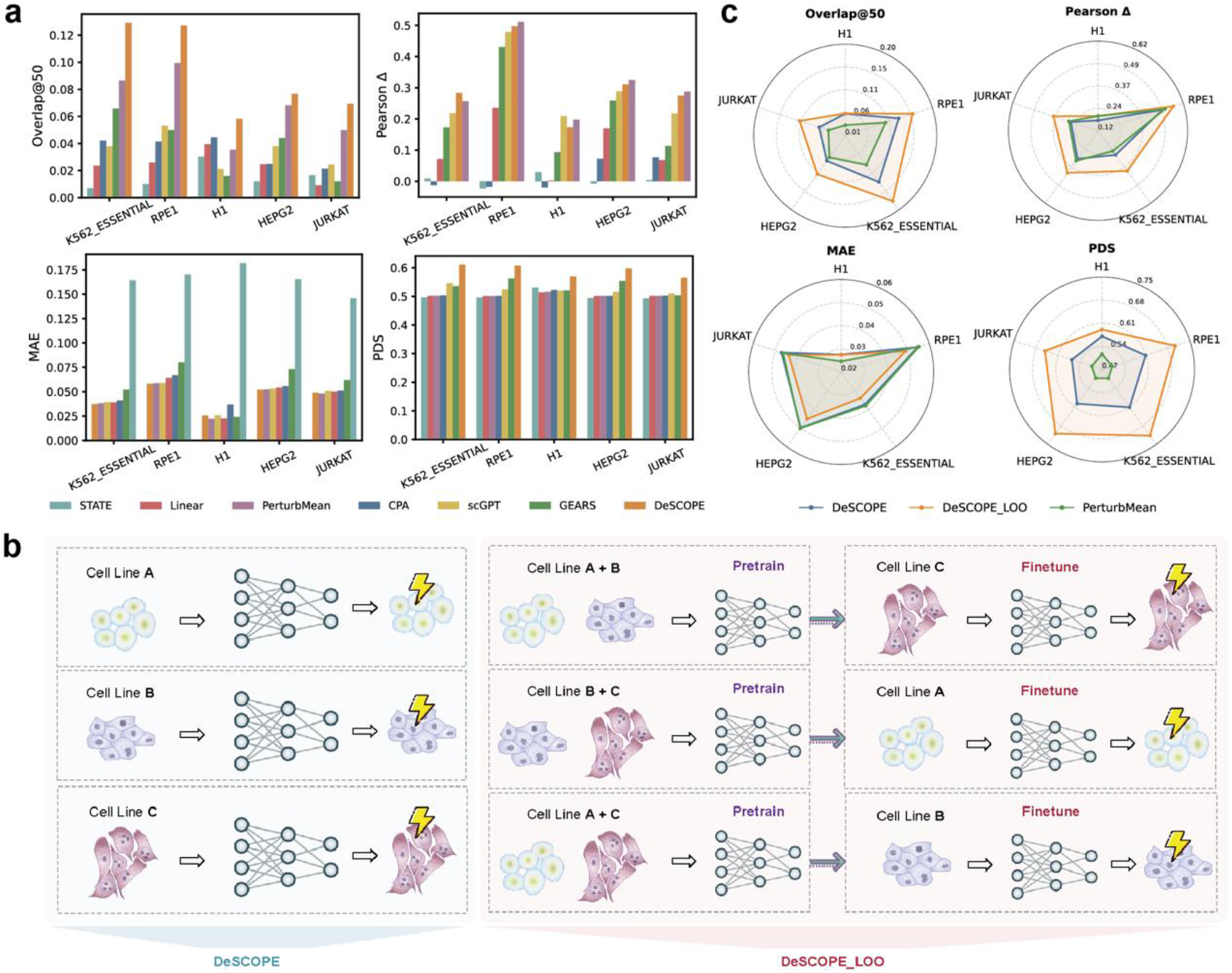
DeSCOPE enables prediction of transcriptomic perturbation responses for unseen genes. **a**, Benchmarking performance on perturbation responses of unseen genes. Overlap@50, overlap ratio of top 50 DE genes; Pearson Δ, Pearson correlation between predicted and true expression changes across all genes; MAE, mean absolute error; PDS, perturbation discrimination score. **b**, Overview of single-dataset and leave-one-out (LOO) transfer learning strategy. **c**, Comparison of DeSCOPE, DeSCOPE_LOO, and PerturbMean on perturbation responses of unseen genes.

However, we observed that DeSCOPE did not consistently outperform the simple baseline PerturbMean on the Pearson Δ metric across several datasets, suggesting that predicting precise expression changes remains challenging when training on a single dataset. Inspired by large language model pretraining strategies and the multi-dataset training paradigm used in STATE, we hypothesized that leveraging perturbation patterns across multiple cell lines could improve predictive performance. We therefore designed a leave-one-out (LOO) transfer learning strategy (Fig. 2b). Specifically, in each experiment, four datasets were first used for initial pretraining to capture shared perturbation-induced transcriptional patterns across cell lines. The pretrained model was then fine-tuned on the remaining dataset using the same training procedure as the single-dataset setting. This strategy enables the model to leverage cross-cell-line perturbation knowledge while adapting to the specific transcriptional characteristics of the target cell line, thereby improving overall prediction performance.

Compared with training on a single dataset, DeSCOPE_LOO consistently improved prediction performance across all evaluation metrics and datasets (Fig. 2c and Extended Data Fig. 2b). Specifically, DeSCOPE_LOO showed consistent improvements across all evaluation metrics, outperforming both the previously superior PerturbMean baseline and the single-dataset DeSCOPE model. Overall, these results demonstrate that the LOO transfer learning strategy enables DeSCOPE to effectively integrate shared perturbation patterns across multiple cell lines, leading to consistently improved and robust predictions of single-gene perturbation responses across diverse cellular contexts.

### Generalization to unseen cell types in transcriptomic perturbation prediction

Cellular responses to genetic perturbations are not conserved; the same perturbation can produce highly heterogeneous responses across different cell types. Therefore, in addition to evaluating predictive models on completely unseen genes, we also assess their generalization ability to entirely unseen cell types. To construct this benchmark, we selected four cell types: K562, HepG2, RPE1, and Jurkat. Due to the substantial computational cost of exploring the full probability space, we simplified the analysis by testing the transferability from a single source cell type (K562) to the three target cell types (HepG2, RPE1, and Jurkat). We built three benchmark datasets that share the same perturbed genes across distinct cell types with available ESM2 embeddings. Specifically, the number of perturbed genes in the datasets is as follows: 1,791 genes for K562 to HepG2, 1,799 genes for K562 to Jurkat, and 1,761 genes for K562 to RPE1. In addition to the baseline methods used in previous sections, we introduced a novel non-learning baseline, DeltaTransfer. This model generates predictions by transferring the perturbation effects of specific genes observed in the source cell type and superimposing them onto the control profiles of the target cell type. We evaluated model performance under both zero-shot and few-shot settings.

In the zero-shot setting, DeSCOPE achieved state-of-the-art (SOTA) performance compared with models such as STATE, scGPT, CPA, GEARS, and Linear. However, it underperformed the non-learning baselines DeltaTransfer and PerturbMean on certain metrics (Extended Data Fig. 3). To further explore model adaptability, we incorporated control cells from the target cell type for fine-tuning, allowing the model to better capture target-specific control-state characteristics. Even in this case, DeltaTransfer remained SOTA, while PerturbMean was outperformed by DeSCOPE on Overlap@50, Overlap@100, and PDS (Fig. 3a). These results indicate that, in zero-shot scenarios, many computational models fail to match the performance of non-learning baselines, further highlighting the non-conservative nature of cellular responses to genetic perturbations across cell types.

**Fig. 3.**
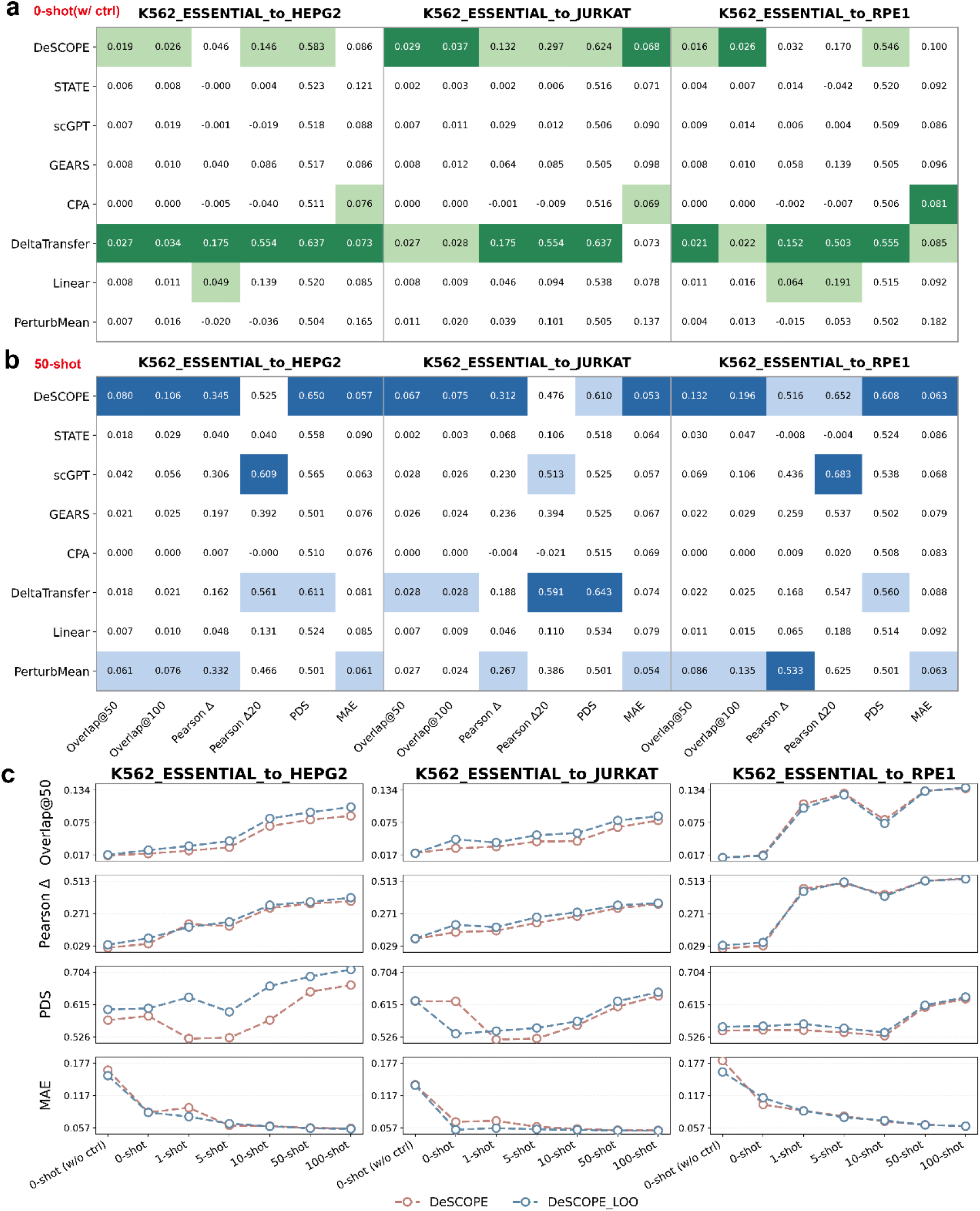
DeSCOPE enables prediction of transcriptomic perturbation responses for unseen celltypes. **a**, Evaluation of perturbation responses of unseen cell types in zero-shot scenarios. **b**, Evaluation of perturbation responses of unseen cell types in few-shot scenarios (50-shot). **c**, Comparison of DeSCOPE and DeSCOPE_LOO on perturbation responses of unseen cell types in four metrics. Overlap@50, overlap ratio of top 50 DE genes; Pearson Δ, Pearson correlation between predicted and true expression changes across all genes; MAE, mean absolute error; PDS, perturbation discrimination score.

To improve cross–cell-type transferability, we explored fine-tuning with limited target-cell-type data. Specifically, we designed a few-shot learning experiment with incrementally increasing numbers of perturbations in the target cell type (1-shot, 5-shot, 10-shot, 50-shot, 100-shot). To ensure fair evaluation, perturbed genes used for training in the target cell type were excluded from the test set. Across all few-shot settings, DeSCOPE achieved near-optimal performance (Fig. 3b and Extended Data Fig. 4), significantly outperforming the non-learning baselines PerturbMean and DeltaTransfer. In contrast, other computational models did not show consistent performance gains. Notably, in the 50-shot setting (Fig. 3b), DeSCOPE improved Pearson Δ20 from 0.14-0.3 to 0.47-0.65 and Overlap@100 from 0.02-0.03 to 0.07-0.2. These results demonstrate DeSCOPE’s ability to rapidly generalize to target cell types with limited data, indicating strong domain adaptation capability.

We also evaluated the effectiveness of the LOO strategy for unseen cell type generalization (Fig. 3c and Extended Data Fig. 5). As the number of perturbations increased, the performance of both DeSCOPE and DeSCOPE_LOO showed an overall upward trend. Importantly, DeSCOPE_LOO provided additional gains over DeSCOPE, especially on the PDS metric, where performance improved by up to 10%. This indicates that by leveraging shared perturbation information across multiple cell lines, the LOO strategy not only improves predictive accuracy in unseen gene transfer tasks but also enhances generalizability in unseen cell type transfer tasks.

### Generalization to different combinatorial perturbation effects prediction

DeSCOPE can be extended to double-gene perturbation tasks. We used the Norman^36^ dataset to evaluate model performance on double-gene perturbations and categorized the settings into three scenarios: Combo_seen0, Combo_seen1, and Combo_seen2, representing the number of perturbed genes observed during the training step (Fig. 4a). We selected the top three models from the Systema^32^ benchmark (MatchMean, PerturbMean, and scGPT) as baselines. DeSCOPE achieves either the second-best or best performance across all scenarios, significantly outperforming both scGPT and PerturbMean (Fig. 4b and Extended Data Fig. 6a).

**Fig. 4.**
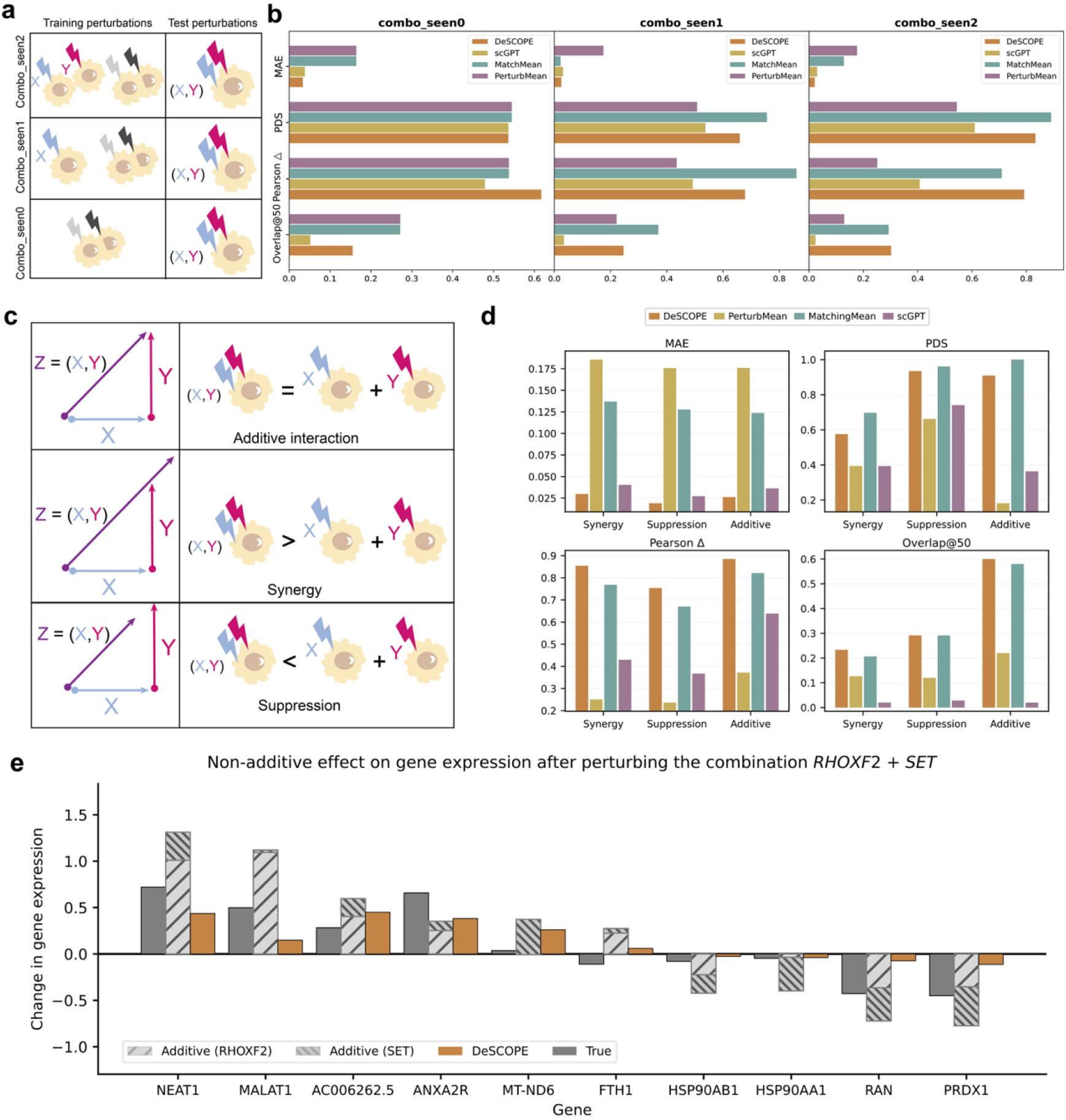
DeSCOPE enables double gene perturbation prediction in different settings. **a**, Train-test data split categories for two-gene perturbations. **b**, Performance comparison under different split categories. **c**, Definitions of the three genetic interaction subtypes. **d**, Performance comparison under different genetic interaction subtypes. **e**, Upon perturbing the RHOXF2 and SET combination, the resulting gene expression changes are displayed. Gray bars denote the actual average expression shift measured after the double perturbation. Hatched gray bars represent the observed changes from each individual single-gene perturbation; their arithmetic sum is what the naive additive model predicts. The orange bar corresponds to the prediction generated by DeSCOPE.

Double-gene perturbations can be classified by their Genetic Interaction (GI) type. In the previous benchmark from GEARS, double-gene perturbations were divided into six categories: Additive, Synergy, Suppression, Neomorphism, Redundancy, and Epistasis. However, since different categories use different evaluation metrics, there is considerable overlap among these six GI types, which inadequately reflects the model’s generalizability across distinct interaction types. To address this issue, we simplified the categories into three major classes: Additive, Synergy, and Suppression (Fig. 4c, see Methods). This approach measures the relationship between the double-gene perturbation effect and the sum of the corresponding two single-gene perturbation effects. This grouping ensures that all double-gene perturbations are divided into three mutually exclusive subsets, thereby better revealing model preferences for different interaction types. We found that under the Combo_seen2 scenario, DeSCOPE achieves the best performance in nearly all metrics, even surpassing MatchMean (Fig. 4d). This suggests that DeSCOPE can effectively leverage known single-perturbation states to reconstruct multi-gene perturbation profiles. In the other scenarios, DeSCOPE also attains near-optimal performance, slightly trailing behind MatchMean (Extended Data Fig. 6b), indicating that DeSCOPE is capable of capturing different GI types. Taking the RHOXF2+SET double-gene perturbation (a Suppressive GI) as an example, we visualized the expression levels of the top ten most affected genes(Fig. 4e). DeSCOPE correctly predicted the direction of change for all these genes. This phenomenon is also observed for other genetic interaction subtypes (Extended Data Fig. 6c).

### Generalization to chromatin accessibility in scATAC-seq perturbation prediction

DeSCOPE can be readily extended to model perturbations across additional modalities. To evaluate its applicability to chromatin accessibility data, we collected five single-gene perturbation scATAC-seq datasets spanning three cell lines (GM12878, K562, and MCF7). The datasets for GM12878, K562, and MCF7 were generated using the Spear-ATAC technology developed by Pierce et al.^37^, while K562_1 and K562_2 were produced in independent batches using the CRISPR-sciATAC method developed by Liscovitch-Brauer et al.^38^. These datasets are characterized by high dimensionality, extreme sparsity, and pervasive zero inflation, which substantially complicate the extraction of genuine biological signals from noise. Following the preprocessing strategy adopted by EpiAgent, raw sequencing peaks were first mapped to a reference vocabulary comprising 1,355,445 candidate cis-regulatory elements (cCREs). The resulting count matrix was then transformed using term frequency-inverse document frequency (TF-IDF) to correct technical biases and highlight informative accessibility signals. Finally, the top 50,000 cCREs ranked by TF-IDF weight were retained as the feature space for downstream modeling. Perturbations were partitioned by target genes, with 75% used for training and the remaining 25% reserved for testing. The perturbed genes in the test set were not observed during training, enabling an evaluation of model generalization in the unseen gene scenario.

Benchmark analyses show that DeSCOPE consistently outperforms EpiAgent across Pearson correlation, Pearson Δ, and MAE (Fig. 5a). Notably, the Pearson Δ for EpiAgent remains below 0.1 in all five datasets, whereas DeSCOPE increases this metric to between 0.30 and 0.65, indicating a substantially improved ability to capture true perturbation effects. To further evaluate the robustness of regulatory direction prediction, we constructed subsets of differential accessibility regions (DA-cCREs) at multiple confidence levels (top 100, 200, 500, 1000, and 2000) and assessed the accuracy of predicting significantly upregulated and downregulated cCREs. As shown in Fig. 5b, DeSCOPE consistently achieves higher directional accuracy than EpiAgent for both upregulated and downregulated regions across all datasets. Moreover, DeSCOPE shows higher median accuracy, a narrower interquartile range, and markedly fewer low-performance outliers. In high-confidence subsets (top100-500 DA-cCREs), directional accuracy remains stable within the range of 0.6-0.8, whereas EpiAgent frequently falls below 0.5. Together, these results indicate that DeSCOPE not only captures global chromatin accessibility patterns more accurately but also more reliably identifies and prioritizes regulatory elements affected by genetic interventions, whether they exhibit activation or repression in accessibilities, thereby providing robust candidates for downstream functional annotation and mechanistic investigation.

**Fig. 5.**
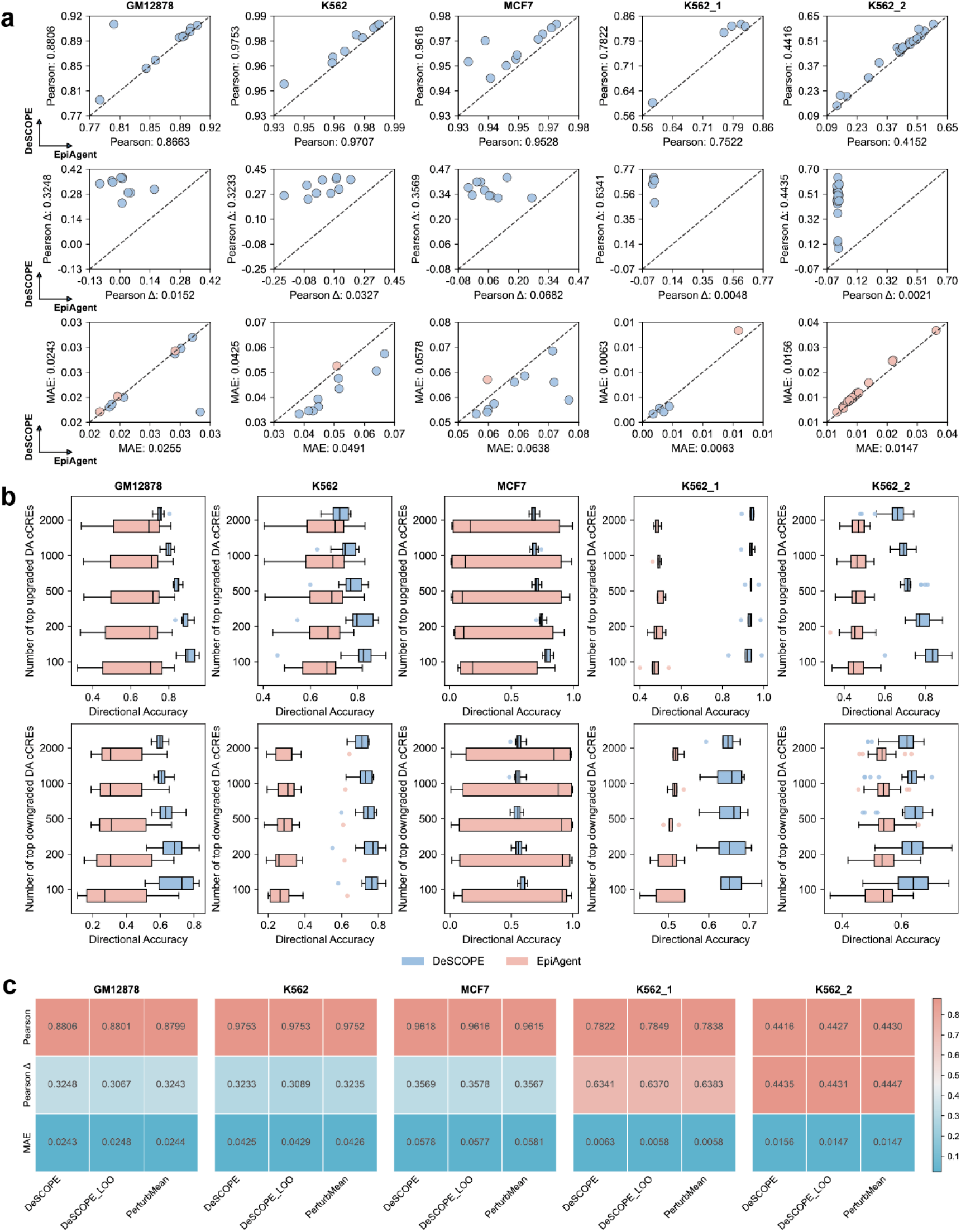
DeSCOPE enables cross-modal prediction of chromatin accessibility perturbations. **a**, Comparison of DeSCOPE and EpiAgent across five single-gene perturbation scATAC-seq datasets (GM12878, K562, MCF7, K562_1, K562_2). Scatter plots show Pearson correlation, Pearson Δ, and MAE between predicted and observed accessibility profiles for held-out perturbation genes; each point represents a single perturbation. **b**, Directional accuracy in predicting significantly up- or downregulated DA-cCREs across different confidence subsets (top 100, 200, 500, 1,000, 2,000). **c**, Performance of DeSCOPE, DeSCOPE_LOO, and PerturbMean across the five datasets.

As an additional evaluation, we adopted the same LOO training strategy used in the scRNA-seq perturbation experiments and compared DeSCOPE with the simple baseline PerturbMean. The results show that DeSCOPE, DeSCOPE_LOO, and PerturbMean achieve comparable performance (Fig. 5c). This observation suggests that simple baselines such as PerturbMean remain highly competitive in scATAC-seq perturbation prediction tasks, indicating that this phenomenon may extend more broadly across perturbation-based omics modeling problems. Importantly, DeSCOPE_LOO does not further improve the performance of DeSCOPE beyond PerturbMean, which may be attributed to two main factors. First, scATAC-seq perturbation datasets typically exhibit extremely high dropout rates (often exceeding 90%), resulting in limited data quality and low signal-to-noise ratios. Second, the top 50,000 cCREs selected from the five datasets show limited consistency across datasets. The overlap between cCRE sets is relatively low, with pairwise intersections generally below 50% (Extended Data Fig. 7).

## Discussion

The emergence of high-throughput single-cell CRISPR screening technologies, combined with the rapid accumulation of large-scale perturbation datasets, has enabled the systematic computational modeling of gene perturbation effects. Nevertheless, most existing approaches still struggle to consistently outperform simple baselines, and their generalization performance remains limited in unseen gene and unseen cell type settings. In this work, we present DeSCOPE, a lightweight and efficient deep learning framework designed to enhance out-of-distribution generalization and modality expansion for single-cell perturbation prediction, with potential implications for small-molecule drug discovery and gene therapy research.

By leveraging gene embeddings derived from the protein language model ESM2 and explicitly decoupling and aligning the latent distributions of perturbed and control cells, DeSCOPE achieves robust performance across multiple benchmark tasks. In particular, in unseen gene prediction, DeSCOPE_LOO is the only method that consistently outperforms the PerturbMean baseline, underscoring the importance of cross-cell-line knowledge transfer for robust extrapolation to previously unobserved perturbation genes. In unseen cell type prediction tasks, DeSCOPE shows a clear advantage in few-shot settings, where fine-tuning with only a small number of perturbation samples from the target cell type yields substantial performance improvements. Furthermore, DeSCOPE demonstrates strong modality generalization, extending beyond transcriptomic readouts to ATAC-based perturbation modeling, providing a flexible framework that can be readily applied to a broad range of single-cell perturbation modalities, including cellular morphology, metabolic states, and other functional phenotypes.

Despite these strengths, DeSCOPE exhibits limited performance in zero-shot unseen cell type prediction settings. This limitation likely arises from the current training strategy, which depends on fine-tuning with perturbation data from the target cell type. When supervision in the target domain is entirely absent, the latent distribution alignment mechanism may be insufficient to accurately calibrate cell-type-specific baseline states, thereby constraining generalization. Future work will investigate the incorporation of cell-type-aware context encoders to explicitly model intrinsic differences between cell types and to provide more informative priors for latent space alignment under unsupervised or weakly supervised conditions. Additionally, integrating larger and more diverse in vitro perturbation datasets may further enhance generalization across genes. Finally, extending DeSCOPE to in vivo perturbation settings represents an important direction for future research, enabling systematic evaluation and optimization of model performance in complex tissue microenvironments. Such extensions may facilitate a deeper understanding of gene function in physiologically relevant contexts and provide more reliable computational support for precision gene therapy and targeted intervention strategies.

## Methods

### Data pre-processing

#### ScRNA-seq pre-processing

For scRNA-seq perturbation data, we first resolved duplicate gene symbols in the raw gene expression count matrix. Within each group of duplicates, only the transcript with the highest mean expression across all cells was retained, ensuring a unique representation for each gene while preserving its most biologically relevant and transcriptionally active signal. After deduplication, counts were normalized by total UMIs per cell and log1p-transformed to correct for differences in sequencing depth and reduce distribution skewness. Due to the high sequencing depth and coverage of the H1 dataset, H1 cells were normalized to 50,000 UMIs per cell, whereas other cell lines were normalized to 10,000 UMIs.

For transfer tasks involving “unseen genes” or “unseen cell types”, training and evaluation often span multiple datasets with inconsistent gene spaces. To ensure cross-dataset compatibility, we first removed duplicate genes within each dataset and then unified the gene feature space by padding the source expression matrix with zero vectors for genes present in the target but absent in the source. Specifically, in the unseen gene task, gene features from all cell lines were mapped onto the gene space of the target cell line for prediction, while the perturbation space remained unchanged. In the unseen cell type task, both gene and perturbation spaces were restricted to their shared intersection across the source and target cell types, and all other training cell lines were projected into this common feature space.

ScATAC-seq pre-processing. For scATAC-seq perturbation data, we adopted the preprocessing pipeline established by EpiAgent. Briefly, the raw cell-by-peak matrix was mapped onto a unified regulatory feature space based on the candidate cis-regulatory element (cCRE) vocabulary defined in EpiAgent. Signals from all peaks overlapping each cCRE were aggregated to construct a cell-by-cCRE matrix. The resulting matrix was then transformed using TF-IDF, after which the top 50,000 cCREs with the highest cumulative TF-IDF scores across all cells were selected as the final feature set for modeling, enriching for cell-type-specific regulatory elements. For multi-dataset joint training, the alignment strategy in the cCRE feature space was kept consistent with that in the gene space.

The aggregated cell-by-cCRE matrix 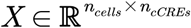, derived from all scATAC-seq data in the Human-scATAC-Corpus, was used to compute TF-IDF scores. Each entry *x*_*ij*_ represents the accessibility count of cCRE *j* in cell *i*. The TF-IDF transformation is defined as:

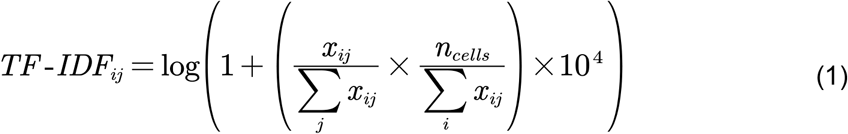

where 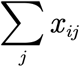 normalizes the accessibility counts within each cell, and 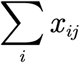 reflects the global abundance of cCRE *j* across the entire corpus.

### Model architecture of DeSCOPE

DeSCOPE is a framework based on cVAE^39^ for modeling cell-state changes induced by gene perturbations. It comprises three key components: a perturbation gene embedding module, a conditional encoder, and a conditional decoder. The embedding module maps high-dimensional representations of perturbation genes into a low-dimensional space, which serves as a conditioning signal for the model. Guided by this gene-level embedding, the conditional encoder processes control and perturbed cells through separate, non-shared networks to produce distinct latent representations. These representations are then used by the conditional decoder to reconstruct and predict cellular features under perturbation.

#### Perturbation gene embedding module

To capture gene-specific heterogeneity in perturbation responses, DeSCOPE leverages ESM2, a protein language model with 15 billion parameters developed by Meta AI, to construct gene embeddings. Specifically, all protein isoform amino acid sequences encoded by a given gene were collected and independently processed by ESM2 to generate contextualized residue-level embeddings. These residue-level embeddings were then averaged across the sequence length to obtain an isoform-level representation. Finally, isoform-level representations corresponding to the same gene were aggregated by mean pooling to produce a unified gene-level embedding. As a result, each gene is represented by a 5,120-dimensional ESM2 embedding vector. For perturbation genes not covered by ESM2, as well as for control cells without any gene perturbation, a zero vector of the same dimensionality was used in our experiment.

The perturbation gene embedding module consists of a linear projection followed by layer normalization. Given an input gene embedding derived from ESM2, denoted as *E*_*gene*_ ∈ ℝ^5120^, the module transforms it into a hidden representation 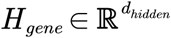 as follows:

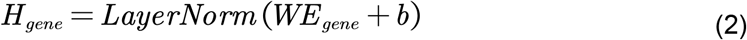

where 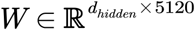 and 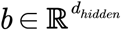 are learnable parameters of the linear layer.

#### Conditional encoder

A key empirical property of single-cell perturbation data is that perturbation effects are often subtle, such that most models struggle to substantially outperform simple mean-based baselines, such as PerturbMean. Accordingly, gene perturbations typically induce only modest shifts in cellular states: perturbed cells tend to remain close to one another in phenotype space and exhibit limited deviation from their corresponding control populations. This behavior is consistent with UMAP visualizations of perturbation data, in which perturbed cells usually show localized displacements from the control distribution rather than forming well-separated clusters. Building on these observations, we depart from the conventional cVAE assumption of a standard normal prior and instead adopt the latent distribution of control cells as the conditional prior. During training, control cells are concatenated with the corresponding gene embedding *H*_*gene*_ and encoded to obtain the prior distribution *P*_*prior*_ (*z*| *X*_*ctrl*,_ *H*_*gene*_). In contrast, perturbed cells are also concatenated with *H*_*gene*_ and encoded by a separate encoder to obtain the posterior distribution *P*_*posterior*_ (*z*| *X*_*pert*,_ *H*_*gene*_). We enforce consistency between the two distributions using a KL-divergence-based regularization term, anchoring perturbed cells to their matched controls in latent space and promoting biologically plausible responses.

The conditional encoder is implemented as an MLP, whose input consists of concatenated cellular features *X* ∈ {*X*_*pert*_, *X*_*ctrl*_} and the gene embedding vector *H*_*gene*_, defined as:

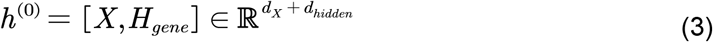

The MLP comprises L stacked layers, each applying a linear transformation followed by LayerNorm, GeLU activation, and dropout:

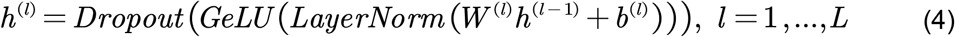

After passing through L layers, the hidden representation *h*^(*L*)^ is projected by a final linear layer to the parameters of a diagonal Gaussian distribution:

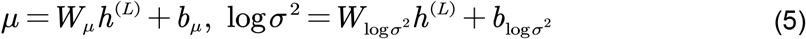

The latent variables *z* ∈ {*z*_*pert*_, *z*_*ctrl*_} is obtained via the reparameterization trick to render the sampling process differentiable:

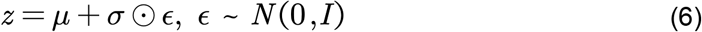

where ⊙ denotes element-wise multiplication.

#### Conditional decoder

In the training phase, the conditional decoder takes as input a latent variable *z*_*pert*_ ∼ *P*_*posterior*_ (*z*|*X*_*pert*_, *H*_*gene*_) sampled from the posterior distribution of perturbed cells, together with the perturbation gene embedding *H*_*gene*_. The concatenated representation is mapped to reconstruct perturbed cellular state. At inference time, since the cellular profiles of perturbed cells are unobserved, the decoder instead operates on a latent variable *z*_*pert*_ ∼ *P*_*posterior*_ (*z* |*X*_*pert*_, *H*_*gene*_) sampled from the conditional prior inferred from control cells. This latent variable is concatenated with the same gene embedding *H*_*gene*_ to predict the corresponding post-perturbation cell state.

The conditional decoder is implemented as an MLP, which predicts the perturbed cellular state as 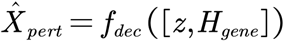. The layer-wise architecture of the decoder mirrors that of the conditional encoder, ensuring consistency in feature representation across the encoding and decoding pathways.

Loss Functions. DeSCOPE is trained by minimizing a composite objective consisting of a reconstruction loss and a latent space regularization term. The reconstruction loss is defined as the mean squared error (MSE) between the observed cellular features *x* ∈ ℝ^*d*^ and their reconstructed outputs 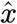:

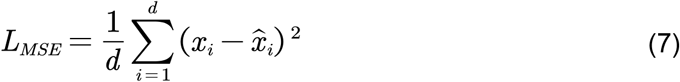

Besides, to regularize the latent space, we further incorporate a Kullback-Leibler (KL) divergence term that penalizes the discrepancy between the posterior distribution inferred from perturbed cells and the prior distribution inferred from control cells, with both distributions conditioned on the same gene embedding:

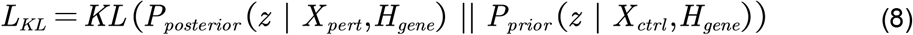

The overall training objective is formulated as a weighted sum of the two terms:

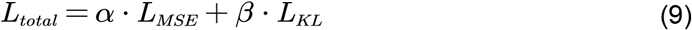

where *α* and *β* are hyperparameters that balance reconstruction accuracy and latent space regularization.

### Implementation details

In our experiments, both the conditional encoder and decoder in DeSCOPE employ an identical architecture consisting of three MLP blocks followed by a final linear projection layer, with a unified hidden dimension of 672, and no dropout was applied during training. Models were optimized using AdamW with a learning rate schedule combining linear warmup and cosine annealing: the learning rate was linearly increased from 0 to a peak value of 1e-4 over the first 10% of training steps, and then decayed to 0 following a cosine schedule. This scheduling strategy improves optimization stability during early training and supports finer convergence in later stages. DeSCOPE was trained for 20 epochs, while DeSCOPE_LOO underwent pretraining and fine-tuning for 20 epochs each. All experiments were conducted with a batch size of 64 on a single GPU.

For the runtime comparison, we used the K562_ESSENTIAL dataset for training. The batch size for all methods was set to 64, and each model was trained for one epoch to standardize the comparison. We compared DeSCOPE with several representative deep learning-based baseline methods that can be trained under comparable computational settings. PerturbMean and Linear were excluded because they do not involve iterative model training and therefore cannot be directly compared in terms of training time and GPU usage under the same settings. STATE was also excluded since its training strategy differs substantially from the other models, making a direct runtime comparison less meaningful.

### Genetic interaction subtypes

In identifying and classifying genetic interactions, we followed the definitions and metrics defined in GEARS. They defined the following types of genetic interactions (GIs): additive, epistatic, neomorphic, redundant, suppressive, and synergistic. However, since different GI types relied on distinct metrics, leading to substantial overlap, we simplified the classification into three categories: additive, suppressive, and synergistic. Here, ‘additive’ interactions are defined as those exhibiting neither synergy nor suppression.

Norman et al. defined metrics (GI scores) for identifying GIs using a linear model of the combinatorial perturbation effect. Let *g*^*i*^ ∈ ℝ^*K*^ be the post-perturbation gene expression vector of a cell *i* with *K* genes. Let *C*_*k*_ be the set of cells under perturbation *k*, where | *C*_*k*_ |= *T*_*k*_. The first step is to compute the average post-perturbation gene expression 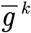 for each of the two combining genes a and b, perturbed singly as well as in combination (a + b):

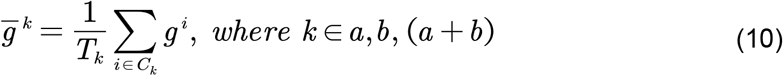

Then the change over mean expression in unperturbed control cells 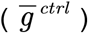 is computed as:

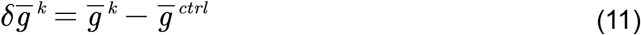

And it is used to fit the following linear model:

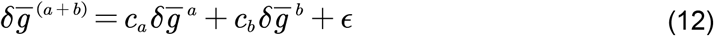

In this model, represents the error term of the model fit. Following the approach of Norman et al., robust regression with a Theil-Sen estimator was used (fitting on 10,000 random subsamples of 1,000 genes each time). Based on the estimated coefficients, the GI scores are defined as Magnitude = 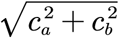. If the Magnitude > 1.15, we consider the genetic interaction type to be synergistic; if the Magnitude < 1.0, we consider it to be suppressive; otherwise, it is additive.

### Evaluation metrics

To comprehensively assess model performance in predicting post-perturbation cell states, we employed a suite of widely used evaluation metrics, including Differential Feature Overlap (Overlap@k), Perturbation Discrimination Score (PDS), Pearson correlation (Pearson), Pearson Δ, and Mean Absolute Error (MAE). Overlap@k measures the extend to which the top-k perturbation-responsive features predicted by a model (e.g., genes or cCREs) overlap with those identified as significantly altered in the ground-truth data. PDS evaluates the ability of a model to discriminate cell states induced by different perturbation genes. Pearson quantifies the overall similarity between predicted and observed post-perturbation molecular profiles, whereas Pearson Δ specifically captures perturbation effects by computing the correlation of changes relative to control conditions. MAE measures the average absolute deviation between predicted and ground-truth values. When top-k differential features were required (e.g., for computing Pearson Δ20), features were ranked in ascending order of false discovery rate (FDR), and the top-k features were selected. All metrics were computed using the open-source evaluation toolkit cell-eval^26^ (https://github.com/ArcInstitute/cell-eval), ensuring consistency and reproducibility of the evaluation pipeline.

## Data availability

All scRNA-seq and scATAC-seq perturbation datasets analysed in this study are publicly available, and the corresponding preprocessing procedures are described in detail in the Methods section. The scRNA-seq perturbation datasets K562_ESSENTIA L and RPE1, generated by Replogle et al.^40^, are available at https://gwps.wi.mit.edu/. The HepG2 and Jurkat datasets, reported by Nadig^41^ et al., are available at Gen e Expression Omnibus (GEO) under accession number GSE264667 (https://www.ncbi.nlm.nih.gov/geo/query/acc.cgi?acc=GSE264667). The H1 dataset was released by t he Arc Institute as part of the 2025 Virtual Cell Challenge^14^ and is available at https://storage.googleapis.com/vcc_data_prod/datasets/state/competition_support_set.zip. T he double-gene perturbation dataset Norman^36^ is available at GEO (GSE133344, https://www.ncbi.nlm.nih.gov/geo/query/acc.cgi?acc=GSE133344). All scATAC-seq perturbation datasets (Pierce et al.^37^, Liscovitch-Brauer et al.^38^), including the cell-by-cCR E matrices, were obtained from the Human-scATAC-Corpus^42^ database and are pub licly available at https://health.tsinghua.edu.cn/human-scatac-corpus/download.php.

## Code availability

DeSCOPE is implemented in the PyTorch framework. Detailed protocols for data preprocessing, model architecture, and training procedures are provided on GitHub (https://github.com/wanglabtongji/DeSCOPE).

## Acknowledgements

This work was supported by the National Key R&D Program of China (2025YFA1805500 to C.W., 2022YFA1106000 to C.W.), the National Natural Science Foundation of China grants (82521002 to C.W.), the Natural Science Foundation of Shanghai (24ZR1492800 to C.W.), Tongji University Medicine-X Interdisciplinary Research Initiative, the Fundamental Research Funds for the Central Universities (22120240435). The authors thank the members of GV20 for helpful discussions on the implementation of DeSCOPE.

## Author contributions

C.W. and X.H. conceptualized the project. P.W., H.W., and Y.L. developed the DeSCOPE algorithm, with P.W. responsible for code implementation and H.W. handling data collection and preprocessing. P.W., H.W., Y.L., and X.Z. conducted benchmarking on various downstream tasks, and collectively drafted the manuscript. X.H. and C.Z provided important suggestions to the framework. All authors provided feedback on the content and figures. C.W. oversaw the overall project and secured funding. All authors have reviewed and approved the final manuscript.

---

**Extended Data Fig. 1.**
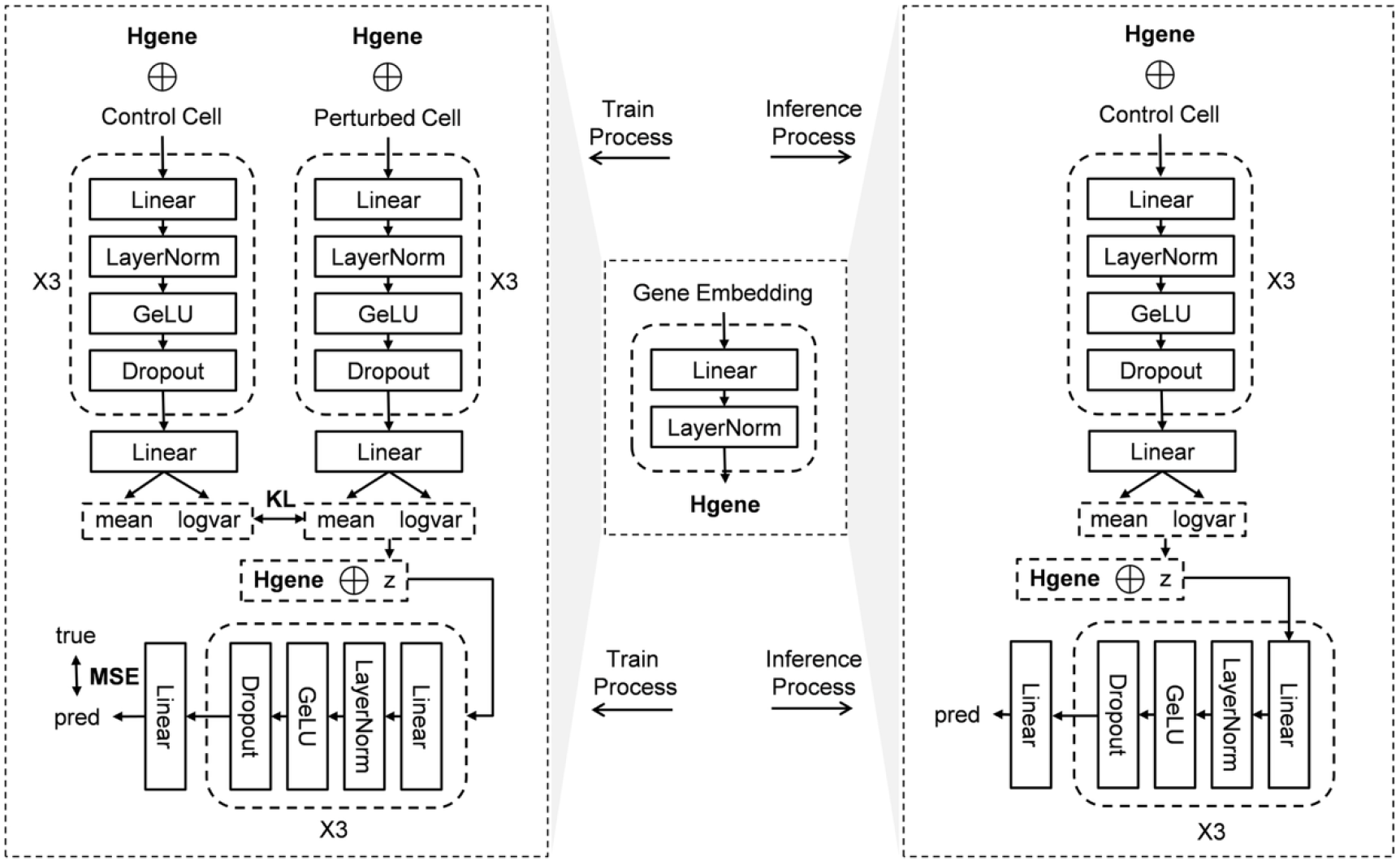
DeSCOPE training and inference pipeline. During training, control and perturbed cell profiles parameterize the prior and posterior distributions, respectively. The decoder reconstructs target expression using latent variables sampled from the posterior. In contrast, during inference where perturbed states are unobserved, the model is conditioned solely on control cells to generate predictions via latent variables sampled from the prior distribution.

**Extended Data Fig. 2.**
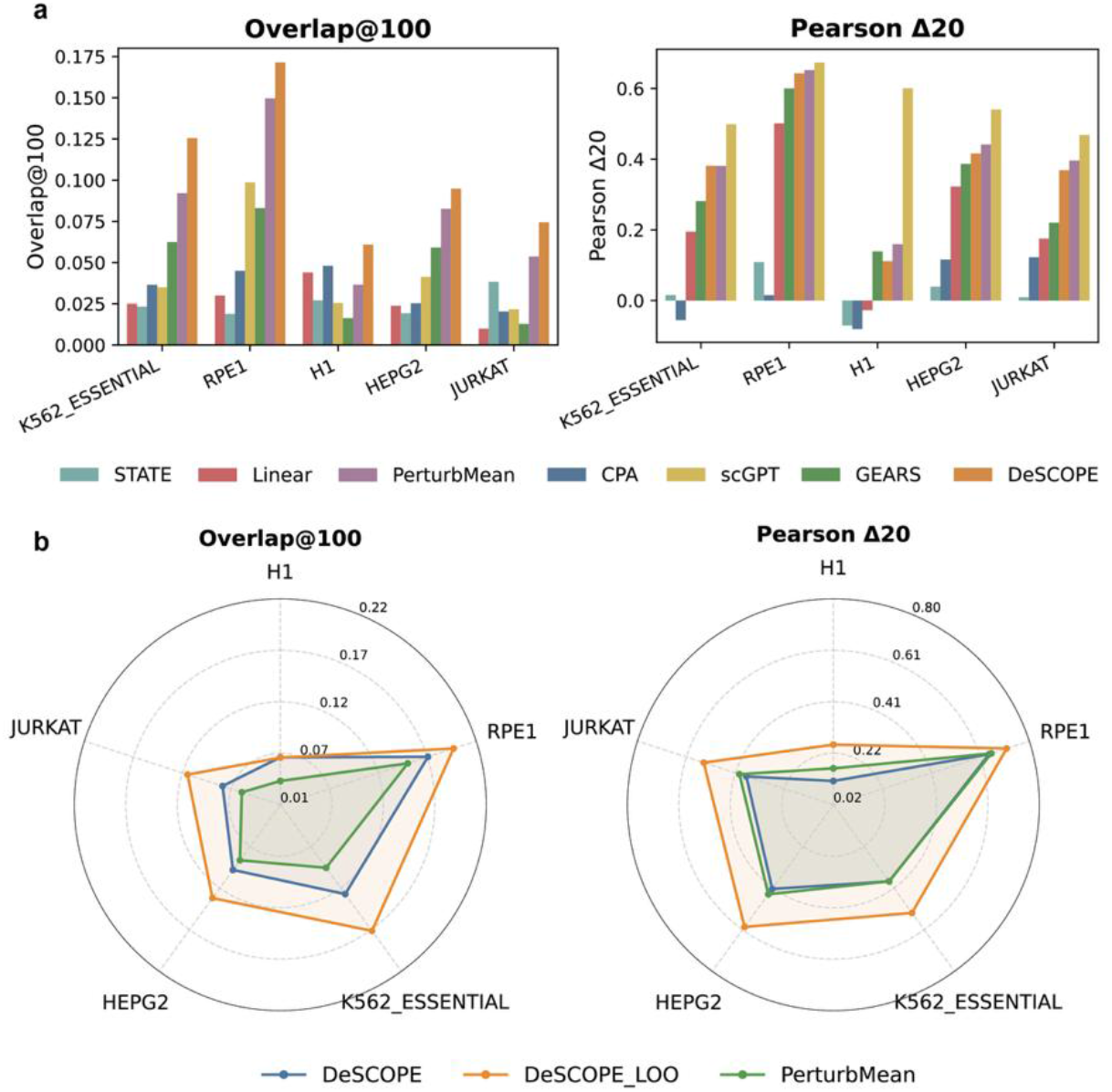
DeSCOPE enables perturbations responses prediction of unseen genes. Benchmarking performance on perturbation responses of unseen genes. Overlap@100, overlap ratio of top 100 DE genes; Pearson Δ20, Pearson correlation between predicted and true expression changes across top 20 DE genes. **b**, Comparison of DeSCOPE, DeSCOPE_LOO and PerturbMean on perturbation responses of unseen genes.

**Extended Data Fig. 3.**
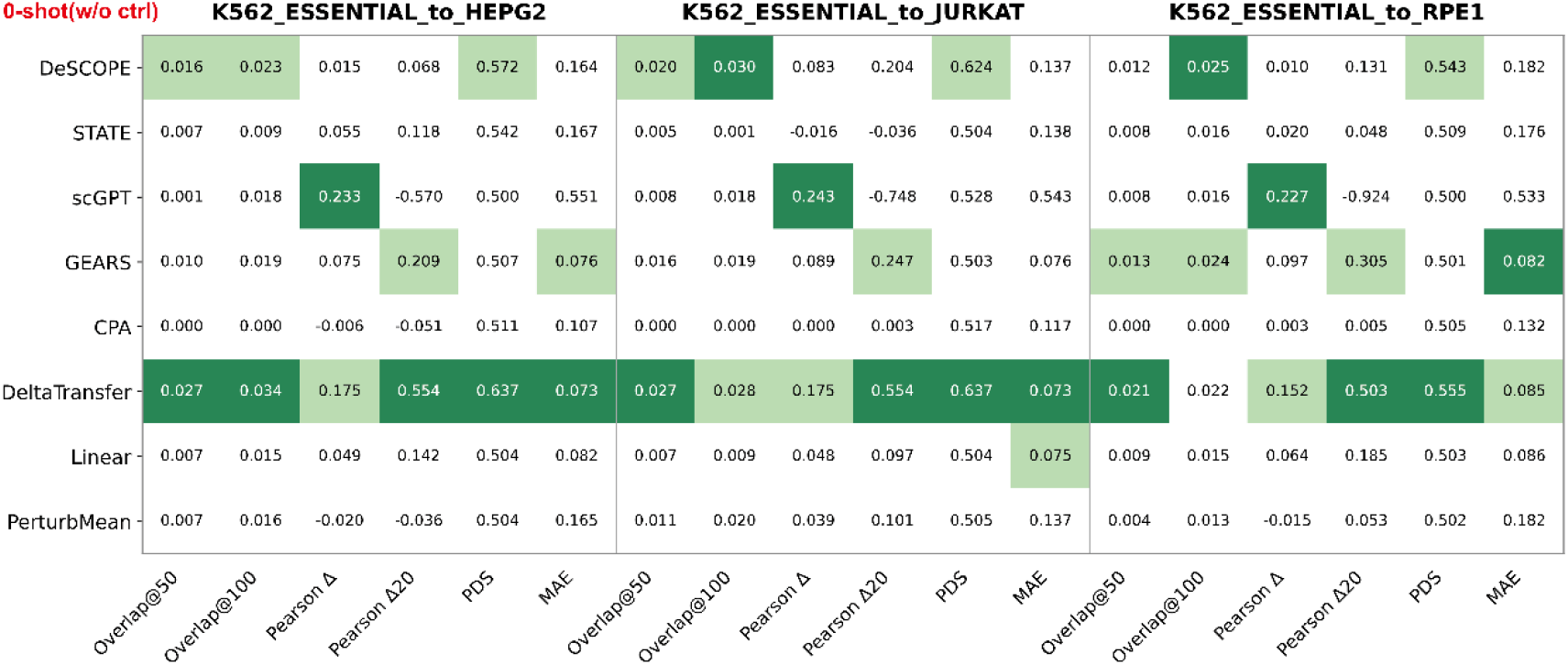
Evaluation of perturbation responses of unseen cell types in zero-shot(w/o ctrl) scenarios.

**Extended Data Fig. 4.**
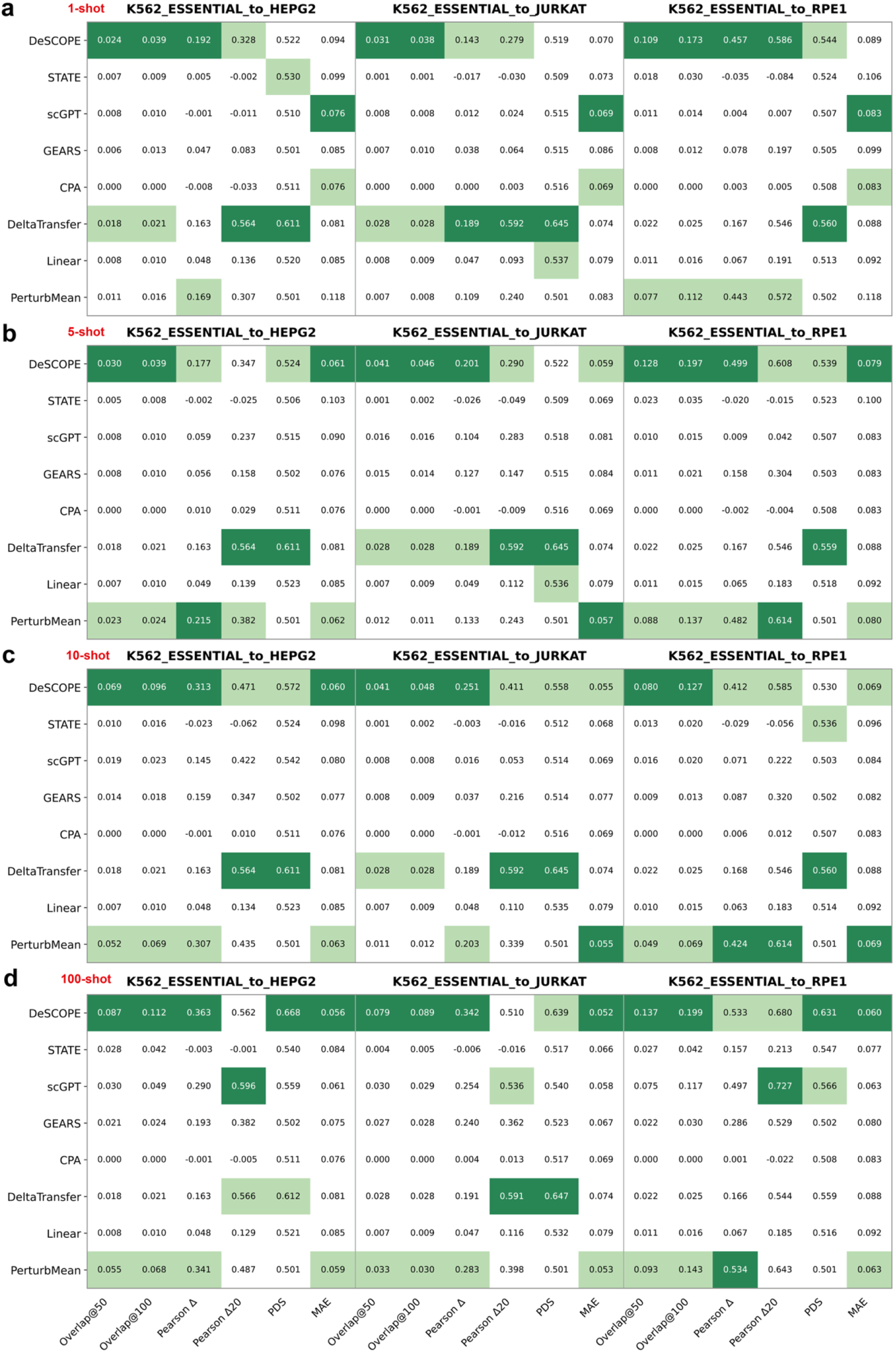
Evaluation of perturbation responses of unseen cell types in few-shot scenarios. **a**, 1-shot scenarios, **b**, 5-shot scenarios, **c**, 10-shot scenarios, **d**, 100-shot scenarios.

**Extended Data Fig. 5.**
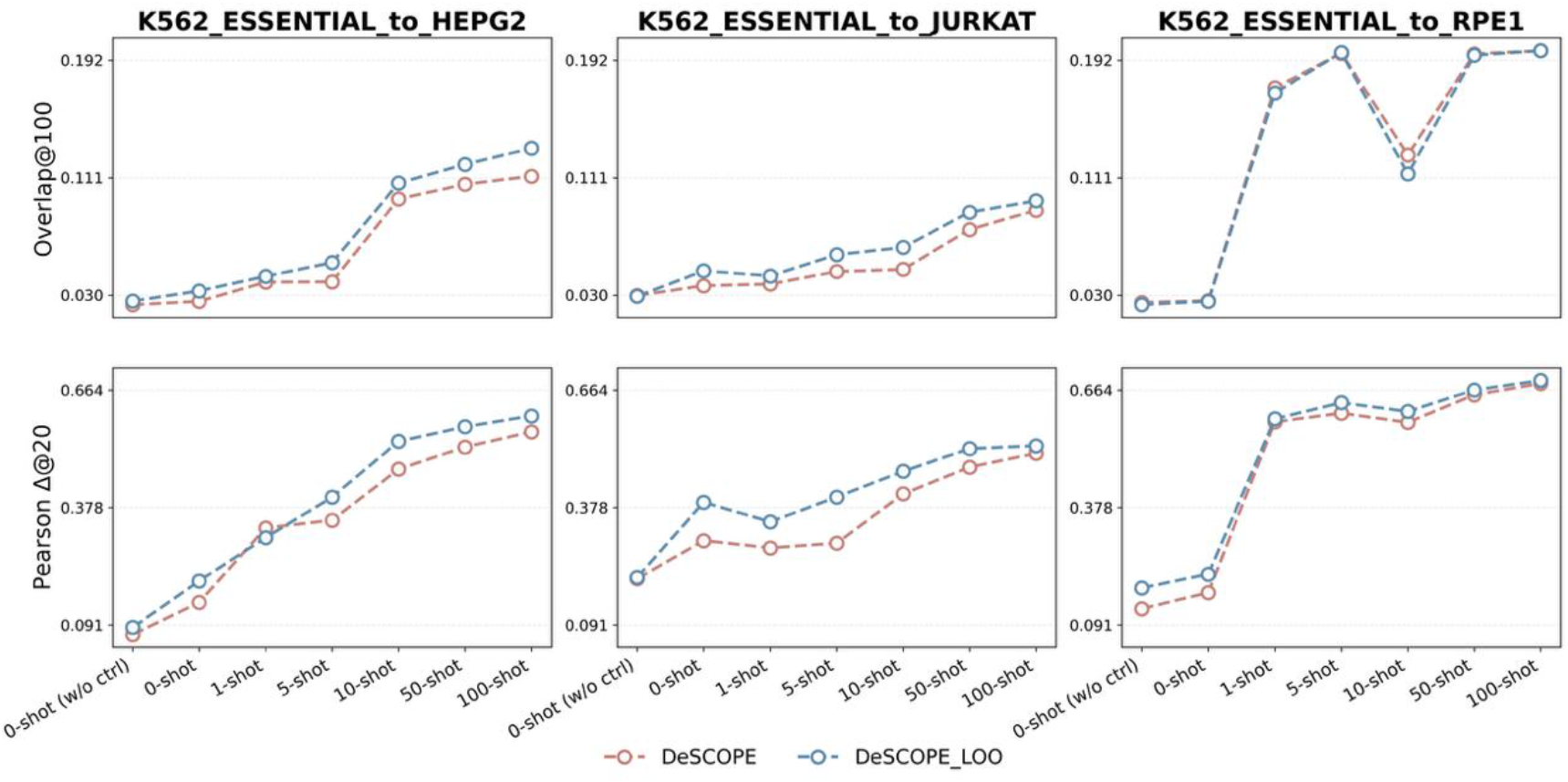
Comparison of DeSCOPE and DeSCOPE_LOO on perturbation responses of unseen cell types in Overlap@100 and Pearson Δ20.

**Extended Data Fig. 6.**
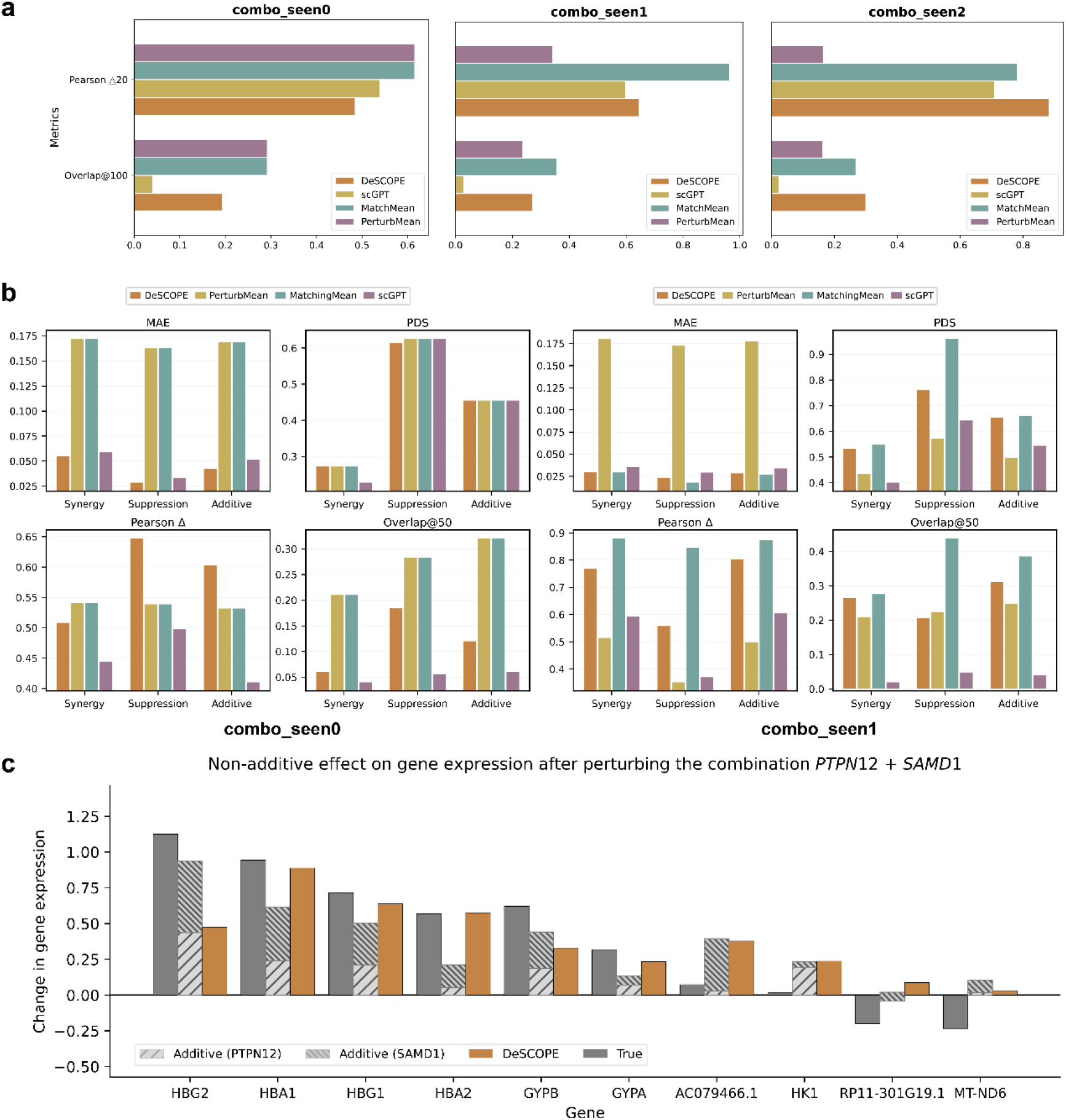
Evaluation of different combinatorial perturbation effects prediction. **a**, Overall performance evaluation in terms of Pearson Δ20 and Overlap@100. **b**, Performance comparison across different GI types under combo_seen0 and combo_seen1 scenarios. **c**, Synergistic effect on gene expression after perturbing the combination PTPN12 + SAMD1. The expression values of the top ten most affected genes are shown.

**Extended Data Fig. 7.**
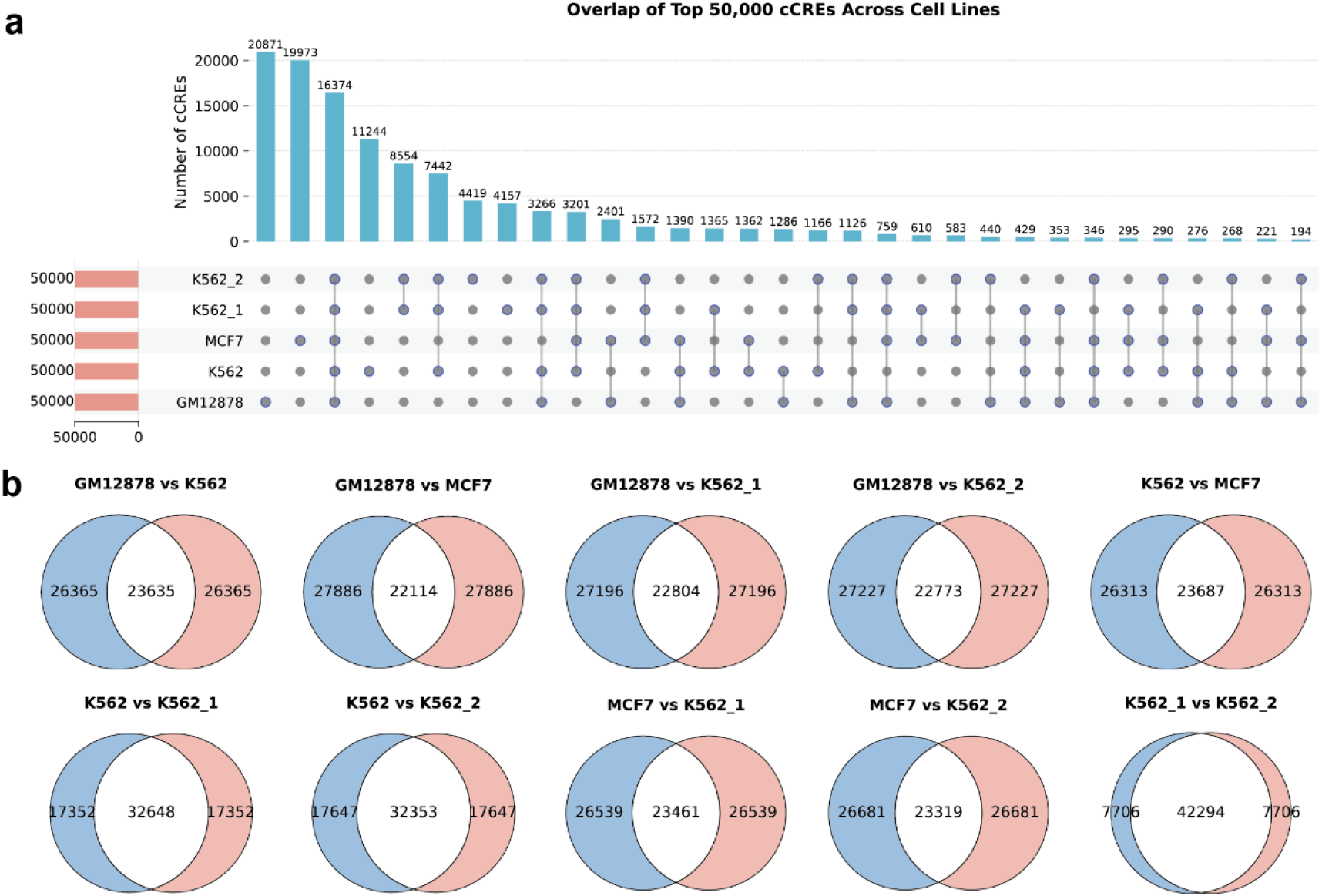
Overlap of top 50,000 cCREs across cell lines. **a**, UpSet plot showing the overlap of the top 50,000 cCREs across five cell lines (GM12878, K562, K562_1, K562_2, MCF7). The bar plot on top indicates the number of cCREs in each intersection, while the connected dots below indicate the specific cell line combinations. The side bars represent the total number of cCREs in each cell line. **b**, Pairwise comparison of top 50,000 cCREs between cell lines using Venn diagrams. Numbers indicate the counts of unique and shared cCREs between each pair of cell lines.

**Extended Data Table. 1.** Comparison of training runtime and gpu memory consumption across different methods on the K562_ESSENTIAL dataset. Experiments were conducted using the K562_ESSENTIAL RNA perturbation dataset. Reported values represent the time required to complete one training epoch with a batch size of 64.

|  | GEARS | scGPT | CPA | DeSCOPE |
| --- | --- | --- | --- | --- |
| Time (s) | 660 | 1464 | 271 | 40 |
| GPU (MB) | 3680 | 25762 | 1034 | 1404 |

